# Neural responses in early, but not late, visual cortex are well predicted by random-weight CNNs with sufficient model complexity

**DOI:** 10.1101/2025.02.05.636721

**Authors:** Amr Farahat, Martin Vinck

## Abstract

Convolutional neural networks (CNNs) were inspired by the organization of the primate visual system, and in turn have become effective models of the visual cortex, allowing for accurate predictions of neural stimulus responses. While training CNNs on brain-relevant object-recognition tasks may be an important pre-requisite to predict brain activity, the CNN’s brain-like architecture alone may already allow for accurate prediction of neural activity. Here, we evaluated the performance of both task-optimized and brain-optimized convolutional neural networks (CNNs) in predicting neural responses across visual cortex, and performed systematic architectural manipulations and comparisons between trained and untrained feature extractors to reveal key structural components influencing model performance. For human and monkey area V1, random-weight CNNs employing the ReLU activation function, combined with either average or max pooling, significantly outperformed other activation functions. Random-weight CNNs matched their trained counterparts in predicting V1 responses. The extent to which V1 responses can be predicted correlated strongly with the neural network’s complexity, which reflects the non-linearity of neural activation functions and pooling operations. However, this correlation between encoding performance and complexity was significantly weaker for higher visual areas that are classically associated with object recognition, such as monkey IT. To test whether this difference between visual areas reflects functional differences, we trained neural network models on both texture discrimination and object recognition tasks. Consistent with our hypothesis, model complexity correlated more strongly with performance on texture discrimination than object recognition. Our findings indicate that random-weight CNNs with sufficient model complexity allow for comparable prediction of V1 activity as trained CNNs, while higher visual areas require precise weight configurations acquired through training via gradient descent.

## Introduction

The development of convolutional neural networks (CNNs) was originally inspired by features of the primate visual system, such as its hierarchical organization [Felleman and Van Essen, 1991, Vezoli et al., 2021] and localized receptive fields (RFs) with repeated feature kernels across space [Hubel et al., 1959, Fukushima et al., 1983, LeCun et al., 1989]. CNNs, and deep neural networks (DNNs) in general, have in turn become effective models of the primate visual ventral stream, allowing for relatively accurate prediction of neural responses to novel, natural stimuli [Cadena et al., 2019, Güçlü and Van Gerven, 2015, Khaligh-Razavi and Kriegeskorte, 2014, Yamins et al., 2014, Zhuang et al., 2021]. The efficacy of task-optimized CNNs in predicting a given brain area’s neural responses (henceforth referred to as “encoding performance”) is thought to depend on several factors, such as the network architecture, the objective function, training dataset and learning rules used for training[Richards et al., 2019, Cichy and Kaiser, 2019, Doerig et al., 2023].

Although earlier work postulated that CNNs can effectively predict neural activity because they are trained on ecologically relevant object recognition tasks [Cadieu et al., 2014, Mehrer et al., 2021], the effect of training may differ substantially between hierarchical levels of the primate ventral stream. It stands to reason that training networks on a diet of natural images with an objective function probing for invariant image classification (e.g. supervised object recognition [Yamins et al., 2014] or contrastive self-supervised learning [Zhuang et al., 2021]) is important for predicting neural activity in higher regions of the primate ventral stream, given their high degree of functional specialization [Cadieu et al., 2014, Rust and DiCarlo, 2010, DiCarlo et al., 2012, Hung et al., 2005]. Indeed, several studies highlight the importance of training objectives and datasets in predicting the activity in higher primate ventral stream regions, while suggesting that differences in architecture between trained CNNs have a smaller influence on explaining responses throughout the primate ventral stream [Storrs et al., 2021, Conwell et al., 2024, Zhuang et al., 2021]. Yet, it is less clear to what extent training CNNs is essential for predicting activity in early visual areas of the primate cortex, which show much less functional specialization and may be involved in a wider range of functions beyond object recognition including scene segmentation [Self et al., 2013], motion processing [Gur and Snodderly, 2007], and salience detection [Li, 2002]. While CNNs trained for object recognition outperform traditional models like linear-nonlinear Poisson models and Gabor filters in predicting macaque V1 responses to natural images [Simoncelli et al., 2004, Willmore et al., 2008, Cadena et al., 2019], this superior prediction may be either due to the CNN’s architecture or the used training objective. A recent study showed that the RF size of neurons in object-recognition trained CNNs was an important determinant of encoding performance, suggesting that the CNN’s architecture does play an important role for predicting V1 activity [Miao and Tong, 2024].

Furthermore, it is plausible that differences between trained and untrained CNNs in predicting neural activity depend strongly on the initial architecture of the CNN with random weights. Recent studies showed that the generalization capacity of DNNs can be attributed to their loss landscapes upon initialization dictated by their architectural design [Chiang et al., 2022, Ramasinghe et al., 2022]. In particular, random-weight DNNs can show strong differences in the complexity of input/output functions dependent on e.g. the non-linear activation function used in the network [Teney et al., 2024]. It is possible that training CNNs steers them towards a certain non-linear complexity matching neural complexity, thereby masking initial differences in architecture, but that a random-weight CNN with an appropriate RF size and non-linear complexity may already allow for accurate prediction of brain activity.

Here, we systematically test whether certain architectural components contribute to CNNs’ ability to encode neural data of early and high visual areas in primates’ brains. Specifically, we constructed CNN models with a linear readout to predict neural data and investigated when training the convolutional filters is necessary for good encoding performance versus only training the linear readouts for an otherwise random-weight CNN.

## Results

### Neural encoding performance of VGG16 model

We analyzed three neural datasets: (1) Firing rates of 166 V1 neurons from two macaques, recorded while the animals passively viewed natural and texture images [Cadena et al., 2019]; (2) activity from 168 multi-unit sites in the IT cortex of two macaque monkeys, passively viewing 3200 grayscale images [Cadieu et al., 2014]; and (3) the fMRI Natural Scenes Dataset (NSD) [Allen et al., 2022], comprising fMRI responses from 8 human subjects viewing thousands of color natural images. Neural activity was predicted by linearly transforming the three-dimensional activation maps of each convolutional layer in the VGG16 model into a one-dimensional vector representing neural activity (either firing rates for macaque datasets or voxel activations for the human fMRI dataset). See supplementary Fig. S1 For a visual illustration of the models. The weights of this linear transformation were fitted on 80% of the dataset and tested on the remaining held-out test set. The “encoding performance” for each VGG16 layer was quantified as the median Pearson correlation between the predicted and actual neural responses across all neuronal sites. Consistent with previous findings, the layers of the ImageNet-trained VGG16 model could predict a substantial amount of variance in neural responses to unseen (i.e. test-set) stimuli in both early and higher visual areas of the macaque and human brain (Fig. 1).

**Figure 1.**
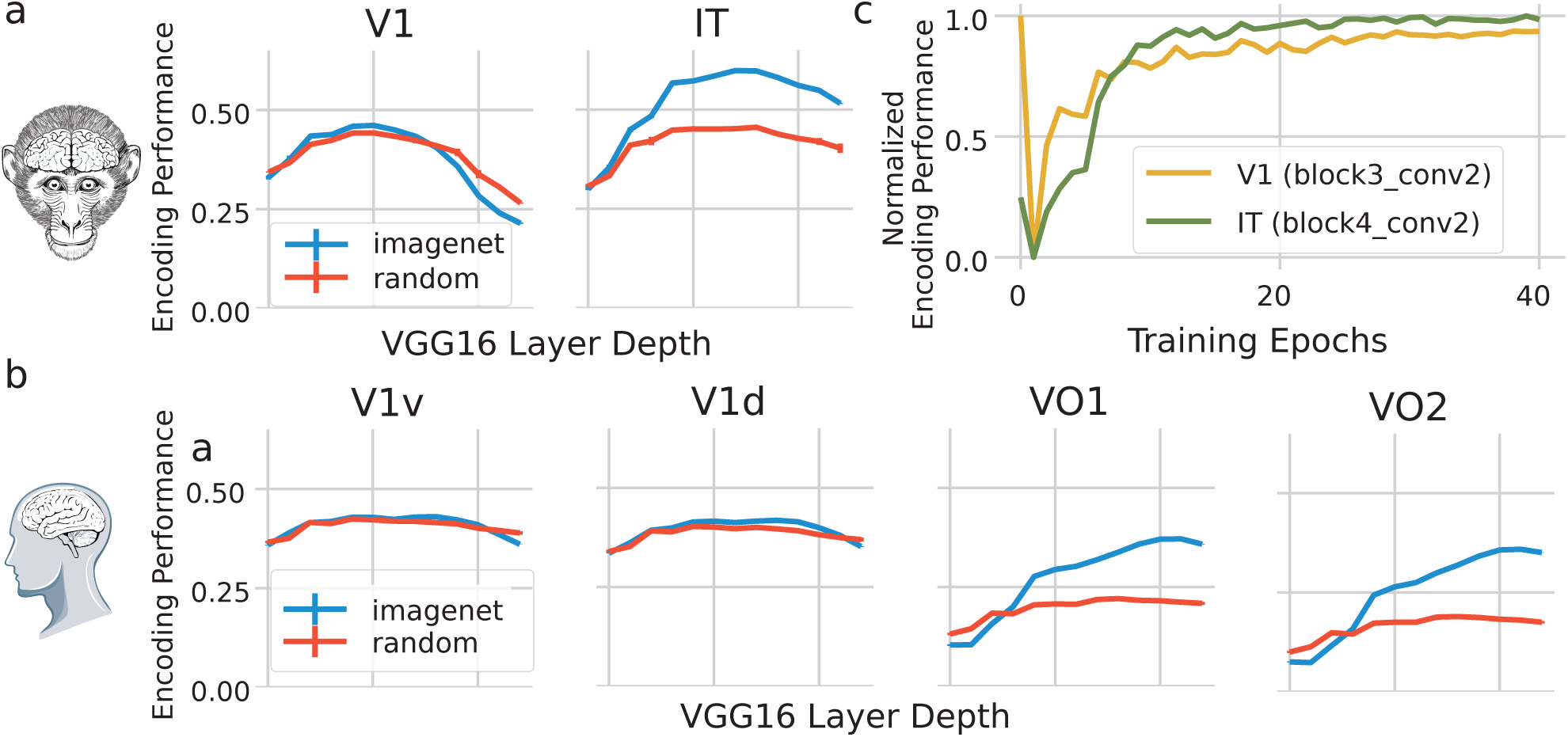
Encoding performance of VGG16 model. **(a)** Encoding performance of a linear readout optimized on top of the representations of the convolutional layers of VGG16 model either upon random initialization (in red) or pretraiend on ImageNet dataset for object recognition (in blue) for two neural datasets recorded from the early visual cortex V1 (left) or the higher visual area IT (right) in macaques. **(b)** same as **a** but for two early visual cortex ROIs (V1v and V1d) or higher visual areas (VO1 and VO2) in huamn fMRI data. **(c)** The normalized encoding performance of the best performing layer for V1 and IT brain areas in macaques tracked over training time on ImageNet dataset from random initialization till full convergence. Each line is the average of 3 training iterations.

Next, we investigated the influence of object recognition training on the encoding performance. When the VGG16 model was initialized with random weights, thereby omitting task training, its ability to predict primate V1 responses showed only minor differences compared to the ImageNet-trained VGG16 model (Fig. 1; difference trained vs. random-weight CNNs: 0.009 for macaque V1, 0.008 for human V1v, 0.012 for human V1d). By contrast, a substantial decrease in encoding performance was observed when predicting responses in higher visual areas using random-weight CNNs (inferotemporal cortex (IT) in macaque and ventral occipital areas (VO1 and VO2) in humans (Fig. 1; difference trained vs. random-weight CNNs: 0.132 for macaque IT, 0.138 for human VO1, and 0.176 for human VO2). The loss in encoding performance for IT was statistically much larger than for V1 (Mann-Whitney U rank test *p* ≪ 0.001). We found qualitatively similar results using other popular convolutional models such as Resnet50 [He et al., 2016], Inception [Szegedy et al., 2015], and DenseNet [Huang et al., 2016] (see supplementary Fig. S3). We furthermore observed only minor differences in V1 encoding performance comparing trained and random-weight CNNs for other measurements of V1 brain activity such as the amplitude of gamma band in local field potentials in an independent data set [Uran et al., 2022] (see supplementary Fig. S4).

To examine the impact of ImageNet training on the encoding performance of VGG16, we quantified the encoding performance across each training epoch, starting with randomly initialized weights and progressing to full convergence (Fig. 1c). Although the network’s performance on object recognition improved monotonically with training, the encoding performance showed a markedly different profile: Starting from the first epoch, the encoding performance for V1 declined notably after the initial training epoch compared to the randomly initialized weights. Hence, training on the object recognition initially decreases the encoding performance, i.e. the ability to predict V1 activity. The encoding performance for V1 only recovered upon reaching full convergence. For IT, however, the encoding performance mostly showed a monotonic increase from the first towards the last training epochs.

Together, these findings indicate that training a CNN architecture (VGG16) on object recognition is not essential for predicting primate V1 activity, as random-weight CNNs demonstrate comparable performance to trained CNNs.

### Simple convolutional models for encoding early and higher visual areas

To further investigate the efficacy of randomly initialized networks in predicting neural responses, we constructed simpler CNN models, systematically changed their architecture and training, and then evaluated their neural encoding performance across various brain regions in macaques and humans. We varied network depth between shallow (2 layers) and deeper (4 layers) architectures, while adjusting convolutional kernel sizes to maintain consistent receptive field sizes across models (see Methods and supplementary Fig. S2). Additionally, we evaluated average and maximum pooling operations and four distinct activation functions (ReLU, ELU, Tanh, and Linear). In this case, we optimized the neural network weights and a linear readout to directly predict neural activity, rather than training on an object recognition task (see Methods), similar to a previous study that developed a shallow neural-network model to predict V1 activity [Du et al., 2024].

For the trained models, we observed that, with the exception of linear networks, all models achieved comparable performance for the V1 area in both macaques and humans (Fig. 2a). Specifically, linear networks employing average pooling exhibited poor encoding performance, while different non-linear activation functions or linear networks utilizing max pooling operations yielded higher and comparable V1 encoding performance. The V1 encoding performance was comparable between shallow and deep CNNs. The performance of the models in predicting responses from higher visual areas (IT, VO1, and VO2) was comparable across non-linear activation functions but improved for deep compared to shallow networks. In sum, when convolutional filters are optimized, the architectural bias of the models is subtle, i.e., the performance difference between different activation functions and pooling mechanisms is almost negligible (except for linear networks with average pooling).

**Figure 2.**
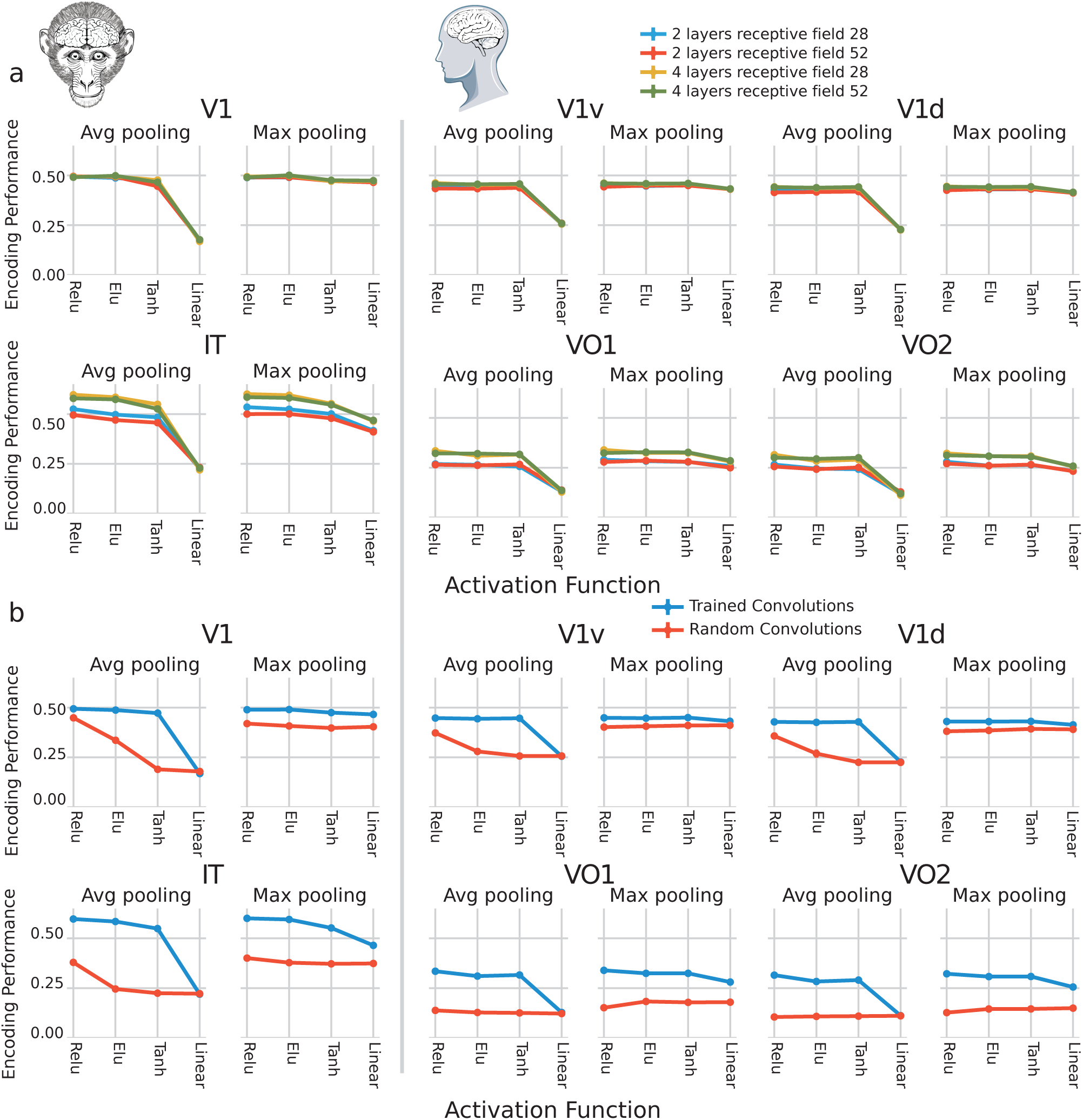
Encoding performance of simple convolutional models on early and higher visual areas of macaques and human brains. **(a)** Encoding performance of shallow (2 layers) and deeper (4 layers) convolutional models each manipulated to have effective receptive field of either 28 or 52 pixels^2^ fully trained on neural data from early (V1, V1v, and V1d) and higher (IT, VO1, and VO2) visual brain areas of macaques and humans. Each model has 8 variants: 2 different pooling strategies (average pooling and maximum pooling) and 4 different activation functions (ReLU, ELU, Tanh, and Linear). **(b)** Encoding performance of the best performing fully trained models from **a** (in blue) – the 2-layer 28 × 28 models for V1, V1v, and V1d brain areas and the 28 × 28 4-layer models for IT, VO1, and VO2 brain areas – and their randomly initialized counterparts (in red). In the latter case, only the linear readout was trained on top of the randomly initialized weights. Each line is the average of 5 training iterations.

We then evaluated the performance of the networks when only the linear readout was optimized, while the convolutional filters were frozen at their randomly initialized weights. For these and subsequent analyses, unless otherwise specified, we focused on shallow 2-layer networks for the early visual cortex and the deeper 4-layer networks for higher visual areas. The comparison of the encoding performance of models with trained convolutional layers to those with randomly initialized weights showed that random ReLU networks approached the performance of their trained counterparts in predicting V1 responses for both macaques and humans (Fig. 2b). The differences between random and trained ReLU networks were 0.045 for the average pooling models and 0.068 for the maximum pooling models. Compared to V1, there was a much greater difference in encoding performance in higher visual areas (IT, VO1, VO2; Fig. 2b).

The differences between random and trained ReLU networks were 0.214 for the average pooling models and 0.196 for the maximum pooling models. In contrast to fully trained networks, networks with randomly initialized convolutional weights showed substantial differences in V1 encoding performance across the activation functions, especially those using average pooling operations (Fig. 2b; *s* = 0.106 for random nonlinear models and *s* = 0.009 for trained nonlinear models).

In summary, random ReLU networks achieved significantly higher encoding performance than other random networks for both early and higher visual cortices. Furthermore, random networks with max pooling operations exhibited substantially higher performance than their counterparts with average pooling, except for ReLU networks, which achieved comparable encoding performance in both scenarios. However, the difference between the best performing random architecture and its trained counterpart was substantially smaller for early visual cortex in comparison to higher visual areas. In conclusion, we identified the ReLU activation function and maximum pooling as key architectural components that significantly contribute to the V1 encoding performance of CNNs. This is evidenced by the comparable performance achieved by randomly initialized models that incorporate these components as compared to their fully trained counterparts.

### Complexity of deep neural networks explains their V1 encoding performance

We showed that random-weight CNNs varied substantially in their encoding performance of brain responses as a function of the non-linear activation function and pooling strategies, especially for early visual cortex (Fig. 2b). However, after having trained the CNNs to predict neural activity, they showed comparable performance across the nonlinear activation functions. Recent studies have shown that random-weight neural networks represent functions of different complexities depending on their architectural components such as their activation functions [Teney et al., 2024, Chiang et al., 2022, Ramasinghe et al., 2022]. We thus wondered whether complexity may explain the encoding performance of random-weight CNNs.

We approximated the function implemented by each output node in each model using a set of Chebyshev polynomials (see Methods). Complexity was then quantified as the average of the polynomial orders, weighted by their coefficients [Teney et al., 2024]. Fig. 3 shows the complexity distribution of output nodes in our models of V1 and IT responses, with both random-weight and trained convolutional kernels (analyses of fMRI models, see supplementary Fig. S5).

**Figure 3.**
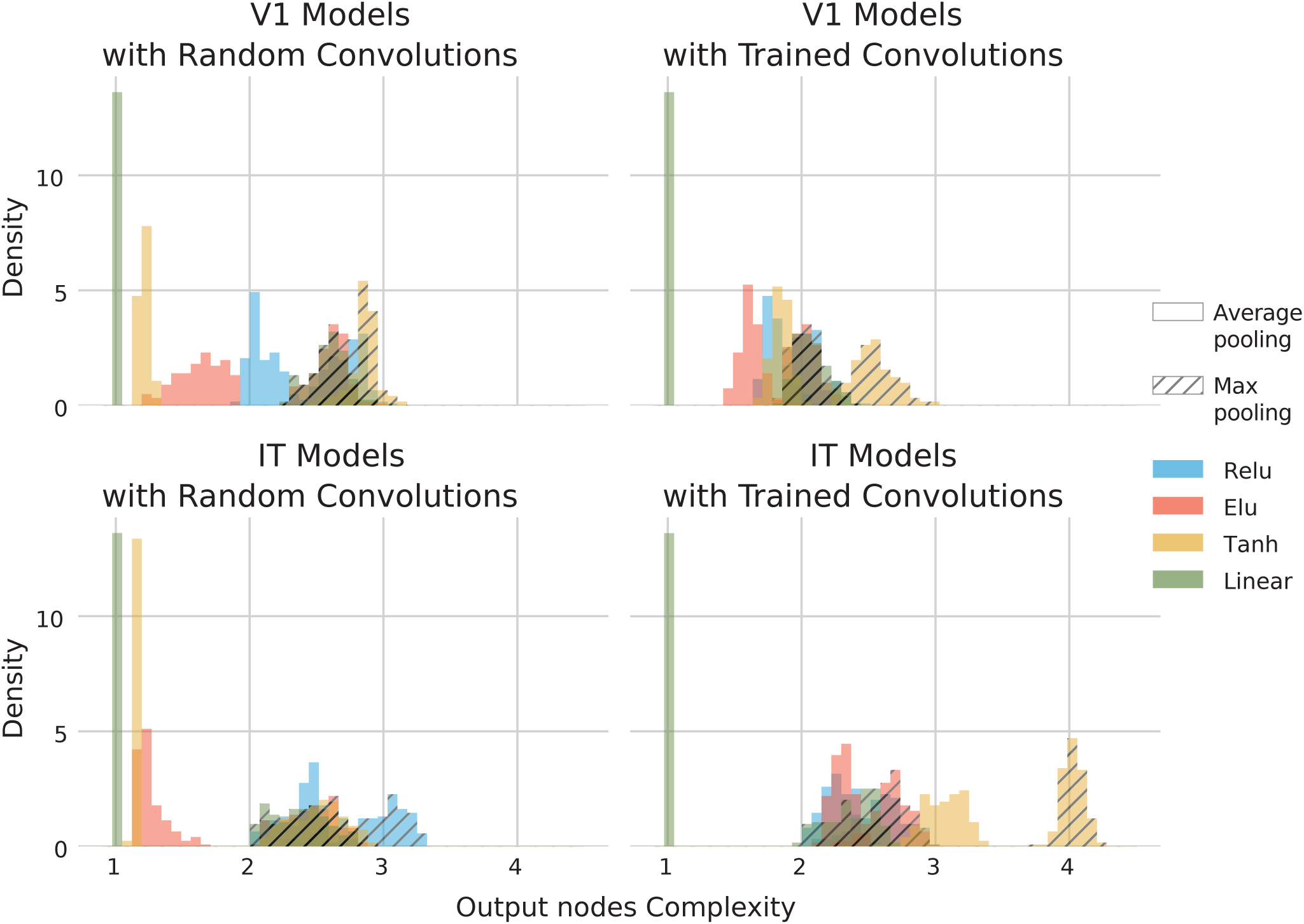
Distributions of encoding models’ complexity across neurons. Each distribution across neurons represent the complexity (see methods) of a certain model configuration (with respect to the pooling strategy and activation function) trained on V1 (upper row) and IT (lower row) data. Only linear readout was trained on top of random convolutional features (left column) or the model was fully trained (right column). Each distribution is the average of 5 training iterations. The corresponding distributions for the human fMRI models are in supplementary Figure S2.

The random-weight CNNs with a low V1 and IT encoding performance (linear, ELU, Tanh with average pooling) generally showed low complexities as compared to their trained counterparts. By contrast, the random-weight CNNs with high V1 and IT encoding performance, such as ReLU and models with max pooling, had higher complexities than the random-weight CNNs with a low encoding performance. Finally, there was a greater overlap among the complexity distributions of models with trained convolutions than those with random convolutions, which is a consequence of the fact that the networks were optimized for the same target function. That is, training CNNs with a different architecture makes their complexity more homogeneous. For V1, random-weight CNNs with a comparable complexity to the trained counterparts also have similar encoding performance, suggesting that model complexity is a main driver of encoding performance. By contrast, for IT, there are major differences in encoding performance between random-weight and trained models despite similar complexity, suggesting that the specific configuration of weights is an additional important factor for IT.

We explored the relationship between model complexities and their encoding performance by plotting the median of the complexity distribution of each model’s output nodes against its encoding performance (Fig. 4). To quantify the relationship between complexity and encoding performance, we fitted a quadratic function. We found a systematic relationship between the median complexity of the models and their encoding performance for V1 in both humans and macaques (Explained variance was 86%, 81%, and 84% for the areas V1, V1v, and V1d respectively). For higher visual areas, the relationship was substantially weaker (Explained variance was 63%, 57%, and 55% for the areas IT, VO1, and VO2 respectively). Specifically, while random-weight models with similar model complexity as trained counterparts had comparable encoding performance for V1, there was major increase in encoding performance for IT. Together, these results suggests that for V1, complexity alone explains encoding performance, while for higher visual areas, the precise configurations of connection weights discovered through gradient descent are crucial for strong encoding performance. To further test the dependence of encoding performance on the precise configuration of weights, we shuffled the weights of all convolutional kernels of all layers of the trained models across all dimensions (space, input channels and output channels), freezing the weights, and subsequently retraining the linear readout. For a fair comparison, the deeper models (4 convolutional layers) were used for both early and higher visual areas. As anticipated, a much stronger decrease in encoding performance was observed for the shuffled models in higher visual areas compared to area V1 for both macaque (Fig. 6a) and human (Fig. 6b) brains.

**Figure 4.**
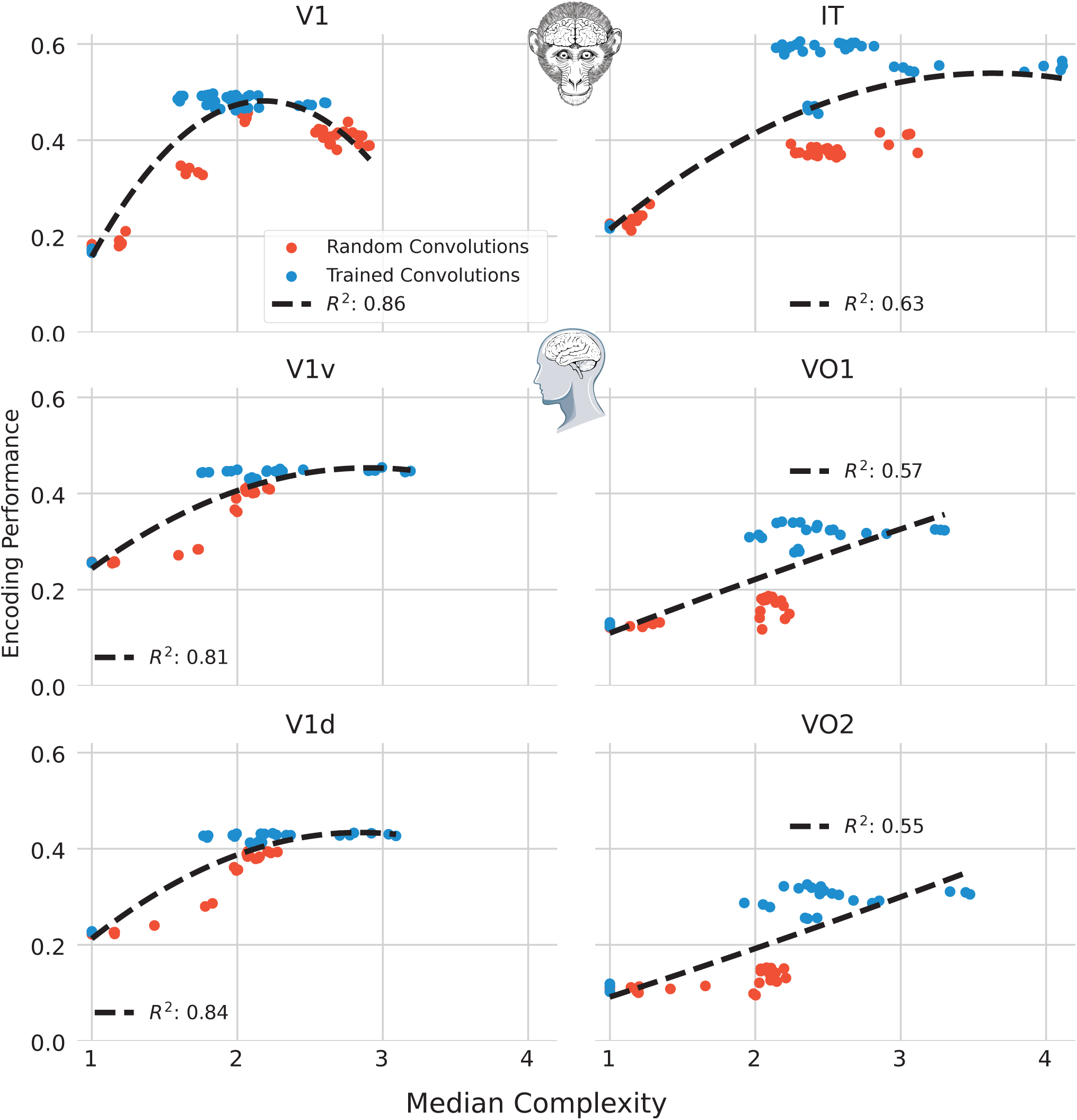
The relationship between the median complexity of the models and their encoding performance. Each panel shows the relationship between the median complexity of the models and their encoding performance for a certain brain area in macaques and humans. Models vary in their activation functions (4), pooling strategy (2) and wether only the readout was trained (in red) or the full model was trained (in blue). Each model configuration is represented 5 times that vary in their random initializations. The dashed black curve represent the fitted quadratic function to the data whose goodness of fit is quantified through *R*^2^ printed in the legend.

### Precise configuration of convolutional weights is critical for object recognition but not texture discrimination

Next, we investigated what kind of visual computations / tasks can be performed by random-weight CNNs, and which tasks are strongly dependent on training. To this end, we created a Texture-MNIST dataset for which two different tasks can be defined (Fig. 5b). Texture-MNIST is a dataset in which every sample is an MNIST digit filled with a texture batch (see Methods). Texture batches are randomly sampled from 10 high-quality texture images. We trained the 4-layer models to predict either the object (digit) identity or the texture patch identity. Similar to the neural data, we either trained only the readout, leaving the convolutional layers frozen at their randomly initialized state, or we trained both the readout and the convolutional layers. We observed that random-weight ReLU networks, with either average or maximum pooling, outperformed all other random-weight networks in predicting the correct identity of the texture class and the digit class (Fig. 5a). However, random-weight ReLU networks achieved almost the same performance as the trained networks on the texture discrimination task, while there was a major difference in performance for digit recognition task.

**Figure 5.**
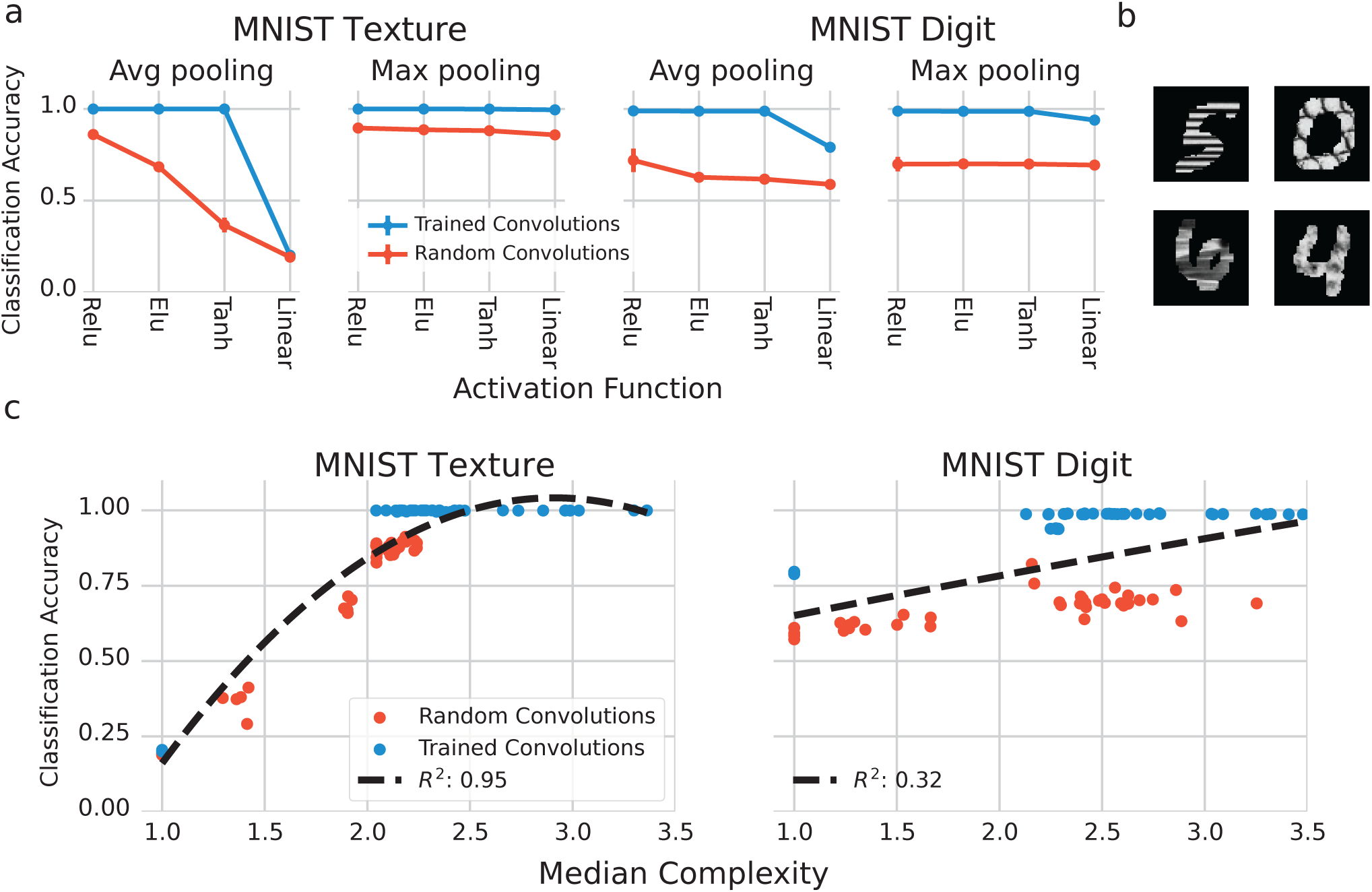
Texture discrimination and digit recognition performance on the Texture-MNIST dataset. **(a)** Performance of our 4-layer models trained on the Texture-MNIST dataset to either predict the texture identity (MNIST Texture) or the digit identity (MNIST Digit). Only the linear classification layer was trained (in red) or the full convolutional model was trained (in blue). Each line is the average of 5 training iterations. **(b)** Examples of the Texture-MNIST dataset. **(c)** The relationship between the median complexity of the mdoels and their texture discrimination accuracy (left) and digit recogntion accuracy (right).

Similar to the neural data, we also investigated the dependence of task performance on the complexity of the models. We found that the complexity of the models showed a very strong relationship with texture discrimination accuracy (explained variance 95%) but not for digit recognition accuracy (explained variance 32%). These findings demonstrate that object recognition performance requires a precise optimization of convolutional kernels, while texture discrimination can already be subserved by random-weight neural networks. We further confirmed this observation by shuffling the weights of the optimized convolutional kernels then retraining the readout. Object recognition performance showed a stronger decrease than the texture discrimination performance (Fig. 6c).

**Figure 6.**
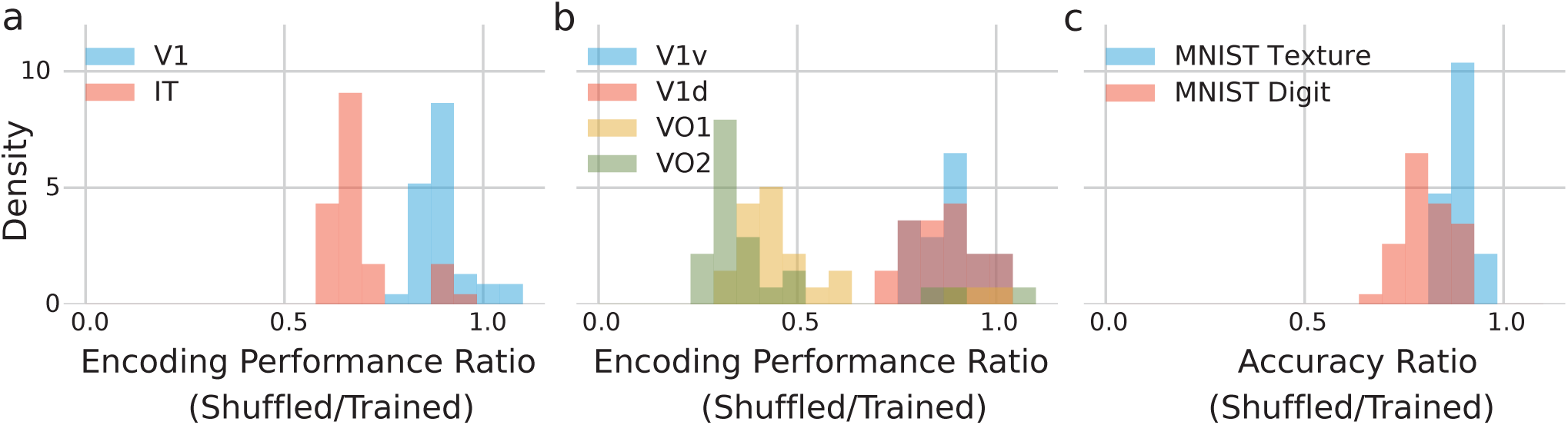
Effect of shuffling the convolutional weights on the neural encoding performance and classification accuracy. For each fully trained model, we shuffled the convolutional weights in the model and retrained the linear readout to predict the neural responses and to classify the texture/digit identity. Then we calculated the ratio between the performance of the model after shuffling and its performance before shuffling. **(a)** The distribution of the encoding performance ratio for all models across different configurations for the macaque brain areas. **(b)** same as **(a)** but for the fMRI human brain areas. **(c)** same as **(a)** but for the models trained for texture discrimination and digit recognition.

### Trained networks develop similar orientation selectivity to V1

We demonstrated that the representations of random-weight CNNs, with an architectural bias that entails sufficient model complexity, suffice for encoding the responses of early visual cortex in macaques and humans. However, it is well-established that V1 neurons exhibit selectivity for certain features, such as the orientation of a bar or grating stimulus [Hubel et al., 1959]. We sought to determine whether random-weight CNNs also possess such feature tuning. To test this, we generated Gabor patches of varying orientations and phases and presented them to the neural networks to assess their orientation selectivity, and then compared the selectivity distributions to experimental V1 data (Fig. 7a). We analyzed the central neurons in the last convolutional layer (i.e. those with a receptive field in the center of the image) of the random-weight neural networks and the networks trained to predict V1 activity. To compute the activation for each orientation, we averaged across all the different phases of the Gabor stimuli. In Fig. 7b, we show the four most orientation-selective neurons in each trained model (for random models, see supplementary Fig. S6a). We then quantified the neurons’ orientation selectivity by calculating the circular variance of their tuning curves. Circular variance is a measure used to quantify the dispersion of data points around a circle, with a lower value indicating more concentrated responses, and thus, higher selectivity [Mazurek et al., 2014] (see Methods). We compared the circular variance distribution of the artificial neurons in the models (Fig. 7c for trained models and supplementary Fig. S6b for random-weight models) with the circular variance distribution of V1 neurons recorded from alert macaque monkeys [Gur et al., 2005]. To this end, we quantified the difference between distributions using the Wasserstein (i.e. Earth Mover) distance, which we term the “V1 deviation score” (lower scores indicate more similarity). Random-weight ReLU networks exhibited the lowest median circular variance among random-weight models (Fig. 7d), i.e. they were the most orientation-selective. Moreover, random-weight ReLU models also demonstrated the lowest V1 deviation score among all random-weight networks (Fig. 7e). Furthermore, training the models on V1 data led to stronger orientation selectivity (i.e. a lower median circular variance) for all the models except the linear ones (Fig. 7d). Moreover, training the models on V1 data also led to lower V1 deviation scores for all of the models except the linear ones (Fig. 7e), with trained ReLU models being the most similar to the V1 orientation selectivity distribution. To examine whether the V1 deviation scores decreased because the networks were specifically trained to predict V1, we also tested the orientation selectivity of the models trained on IT data. We found that IT trained models were less orientation-selective than V1 trained models (supplementary Fig. S7a), and had higher V1 deviation scores (supplementary Fig. S7b).

**Figure 7.**
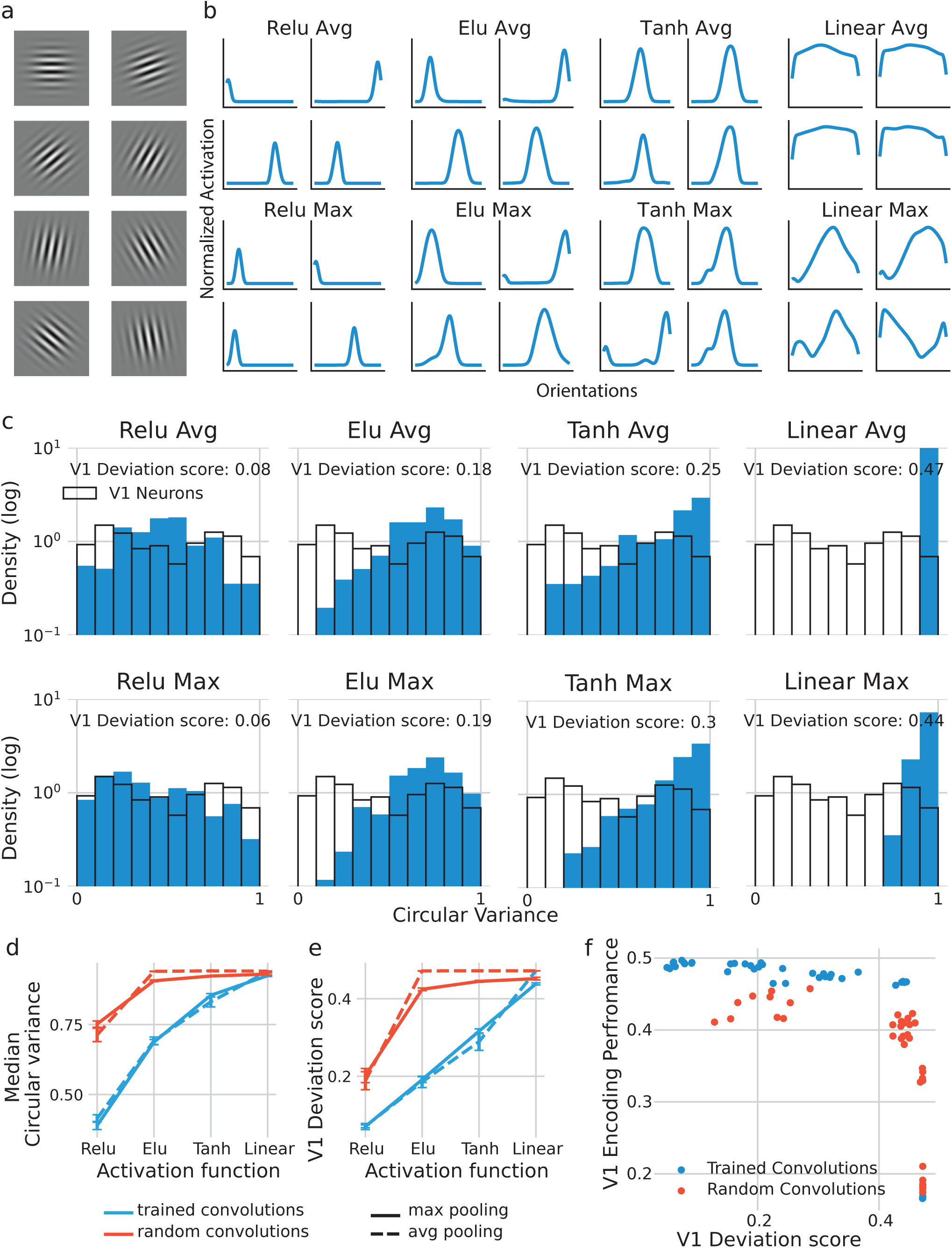
Orientation selectivity of random and trained models. **(a)** Examples of the Gabor patches that were represented to the models. **(b)** Tuning curves of the most orientation selective artificial neuron in the last convolutional layer of each model configuration trained on the V1 data. For the randomly initialized models see supplementary figure S2a. **(c)** Distributions of the circular variance of the artificial neurons of the last convolutional layer of each model configuration trained on the V1 data. For the randomly initialized models see supplementary figure S2b. **(d)** Median circular variance of the artificial neurons of the last convolutional layer of random and V1 trained models of different configurations averaged across 5 iterations. Error bars are the standard deviation. **(e)** V1 deviation scores of the random and V1 trained models averaged across 5 iterations. Error bars are the standard deviation. **(f)** The relationship between V1 deviation score and the V1 encoding performance for all the models that varied in their activation functions, pooling strategies, and their random initializations. Spearman correlation = −0.87, *p* ≪ 0.001.

To examine if networks with more similar orientation tuning to V1 also better predicted V1 activity (on another dataset), we examined the relationship between both variables across random and trained models (Fig. 7f). We found a strong negative monotonic relationship (Spearman correlation = −0.87) between the V1 deviation score and V1 encoding performance, suggesting that the more similar the orientation selectivity of the models is to that of V1 neurons, the better the models are at predicting V1 responses (on another dataset). However, it is also worth noting that although trained networks demonstrated small variation in their V1 encoding performance, they showed high variability in their V1 deviation score (Fig. 7f), suggesting that having similar orientation selecting to V1 neurons is not a strong requirement for the model to be able to effectively encode V1 responses.

## Discussion

We investigated the factors contributing to the success of CNNs in predicting responses within the visual cortex in both macaques and humans. We demonstrate that, unlike higher visual areas, accurate prediction of early visual cortex responses does not necessarily depend on the optimization of convolutional kernels via training. Instead, architectural components, such as pooling strategies and activation functions, play a crucial role. Specifically, we found that the ReLU activation function and maximum pooling were the most critical components for achieving high encoding performance for early visual cortex, even in the absence of optimization of the convolutional kernels based on a task or neural data. Furthermore, we observed that random-weight CNNs exhibited substantial variability in the complexity of the functions they represented, depending on their architectural components. Notably, the CNN’s complexity explained significantly more variance in the encoding performance in early visual cortex compared to higher visual areas. These findings held true across electrophysiological data from macaques and fMRI data from humans. Additionally, model complexity explained significantly more variance in performance on a texture discrimination task than on a digit (object) recognition task using the same dataset. This suggests that precise configuration of convolutional kernels is more essential for object recognition and, consequently, more critical for predicting responses in higher visual areas. Importantly, our results indicate that training the full convolutional models masked the effect of architectural bias, as the variability in encoding performance across different model configurations decreased significantly after training. However, when employing an alternative metric for aligning models with experimental V1 data – namely, the circular variance distribution of the model’s artificial neurons as a proxy for their orientation selectivity – a different picture emerged: Trained models exhibited significant variability in their alignment with V1 orientation selectivity, despite displaying low variability in their V1 encoding performance. Overall, our findings indicate that random-weight CNNs with sufficient model complexity allow for comparable prediction of V1 activity as trained CNNs, while higher visual areas require precise weight configurations acquired through training via gradient descent.

Our work highlights the importance of random-weight controls to reveal architectural bias. According to the deep learning framework for neuroscience [Richards et al., 2019], various deep neural network models, trained under specific constraints regarding their architecture, objective function, and training data, can serve as tests for specific hypotheses about brain function [Zhuang et al., 2021, Dwivedi et al., 2021, Doerig et al., 2023, Cichy and Kaiser, 2019]. The degree to which these models’ representations of complex natural stimuli align with brain responses can provide evidence for certain hypotheses, such as the necessity of recurrent connections for cortical information processing [Kietzmann et al., 2019, Kubilius et al., 2019]. However, it has been shown that multiple models with different architectures, trained similarly, or multiple models with the same architecture, trained differently, can achieve similar performance in predicting neural data [Storrs et al., 2021, Conwell et al., 2024]. Further investigations revealed that task optimization increased the effective dimensionality [Del Giudice, 2021] of CNNs representations. This increase in dimensionality correlated with their encoding performance of higher visual brain areas [Elmoznino and Bonner, 2024]. A recent study showed that substantially scaling the dimensionality of random CNNs, but not transformers or fully connected networks, led to a comparable encoding performance of higher visual areas to ImageNet-trained AlexNet model suggesting a strong contribution of architecture bias to DNN encoding performance of neural data [Kazemian et al., 2024]. In the present work we showed that scaling the complexity of random CNNs, in the low-dimensional regime, leads to a comparable encoding performance for lower, but not higher visual areas. Moreover, it has been shown that layers of task-optimized VGG19 and Alexnet models with matching receptive field sizes yielded similar V1 encoding performance despite having different depths i.e. different numbers of nonlinear transformations suggesting the significance of receptive field size in predicting V1 data [Miao and Tong, 2024]. These findings underscore the necessity of implementing appropriate controls when employing task-optimized or brain-optimized DNNs for predicting neural activity. Specifically, the use of random-weight models with variable architectural components is crucial to reveal the architectural biases of neural encoding models.

Beyond the hypothesized neural factors influencing the neural encoding performance of DNNs, their success in predicting brain data could originate from non-neural, biologically implausible design choices implemented by researchers. For instance, the emergence of grid-like representations in DNNs optimized for path integration has been shown to depend on specifically designed readout mechanisms [Schaeffer et al., 2022]. Moreover, numerous studies have demonstrated that the measured similarity between models and the brain can be highly dependent on the chosen similarity metric [Soni et al., 2024, Davari et al., 2022]. Specifically, neural predictability scores based on linear regression can be heavily influenced by the inductive biases of linear regression, the dimensionality of the model representations, or the ratio of the number of stimuli in the benchmarking dataset to the dimensionality of the model representations [Canatar et al., 2024, Schaeffer et al., 2024, Bowers et al., 2023, Elmoznino and Bonner, 2024]. When we assessed the brain alignment of V1 models using two independent scores: encoding performance on one dataset and V1 deviation score on another dataset, we found large difference in the variability of both scores despite correlating with each other. Therefore, consistent with the existing literature, our findings highlight the importance of moving beyond single metrics of model-brain alignment. Instead, our findings highlight the importance of a multidimensional model assessment approach that enables the dissection of the similarities and differences between computational models and the brain [Wichmann and Geirhos, 2023, Biscione et al., 2024, Jacob et al., 2021, Rajesh et al., 2024]. This holds true in particular considering that many of these similarity measures are correlational [Bowers et al., 2023], and models with high prediction scores can still operate in qualitatively different ways than the brain [Geirhos et al., 2019, Geirhos et al., 2018, Baker et al., 2018, Farahat et al., 2023, Wichmann and Geirhos, 2023]. This multidimensional approach will facilitate more targeted model improvement and informed hypothesis generation in future research.

By systematically manipulating the architectural components of the models and comparing the performance of randomly initialized models with their fully trained counterparts, we identified the essential components that underpin neural encoding, raising the question to what extent they mimic the architecture of visual cortex. The ReLU activation function emerged as a key factor in generating visual representations that supported the most efficient encoding performance, considering the number of trainable parameters. The ReLU activation function was introduced to DNNs as a more biologically plausible alternative to Sigmoid and Tanh functions, given its one-sided nature (outputting zero for negative inputs) and its promotion of sparse representations [Glorot et al., 2011, Attwell and Laughlin, 2001, Douglas et al., 1995]. Indeed, ReLU networks, even with random convolutions, not only exhibited the best encoding performance for V1 but also displayed the smallest distance to the orientation selectivity distribution of V1 neurons. Importantly, while models fully fitted to predict V1 data, with ReLU, ELU, or Tanh activation functions, exhibited similar encoding performance, they still displayed substantial variability in their similarity to V1 orientation selectivity, with ReLU networks being the most V1-like. This result demonstrates that the combined application of multiple model assessment metrics and systematic architectural manipulations enables the identification of key, potentially biologically plausible architectural components that contribute significantly to neural encoding performance. Furthermore, the fact that non-linearities with sufficient model complexity are a major factor in predicting neural activity fits with the general idea that non-linearities are a central component of cortical inter-areal interactions beyond mere linear information transmission [Vinck et al., 2023, DiCarlo et al., 2012].

Beyond examining the encoding performance and tuning properties of the models’ representations, it is imperative to understand the computational advantages of models’ representations in supporting visual tasks. We demonstrated that random-weight CNN representations, which were sufficient for predicting early visual cortex responses, performed well in discriminating between texture families compared to their trained counterparts. Conversely, these random-weight representations were considerably worse than fully trained models in invariantly classifying the identity of digits within images. Studies have shown that V1 activity, in particular superficial cortex, exhibits selectivity for texture statistics, albeit less pronounced than in a higher visual area, LM [Bolaños et al., 2024, Ziemba et al., 2019]. In humans, texture discrimination task-learning has been shown to induce local changes within the early visual cortex without requiring the recruitment of higher visual areas [Schwartz et al., 2002]. Additionally, a decoder trained on macaque V1 population activity elicited by texture samples could discriminate between 15 different texture families [Ziemba et al., 2016]. In contrast, several studies have demonstrated that IT neurons possess the tolerance to identity-preserving transformations that is essential for object recognition [Rust and DiCarlo, 2010, Hung et al., 2005].

The efficacy of random features in machine learning has been well-documented, often rivaling hand-crafted or even learned features across various learning tasks [Rahimi and Recht, 2008a, Gallicchio and Scardapane, 2020, Scardapane and Wang, 2017]. Random-weight CNNs were shown to be frequency-selective and translation-invariant which explains their superior performance over random non-convolutional networks on image classification tasks [Saxe et al., 2011]. Moreover, only training a small fraction of the convolutional weights or only training the batch normalization layers in random-weight CNNs led to object recognition performance competitive with their trained counterparts [Rosenfeld and Tsotsos, 2019, Frankle et al., 2021]. Furthermore, the structure of a random generator CNN can capture significant low-level image statistics even without any learning. This inherent structure acts as a prior, making random-weight CNNs useful for various image processing tasks such as image restoration, denoising, inpainting and super-resolution [Ulyanov et al., 2018]. These findings emphasize the significant contribution of convolution and pooling operations, independent of learning, in visual processing tasks. Consequently, it is plausible that random features with the right convolutional architectural bias could effectively model the representations found in V1, considering the diverse range of visual tasks that V1 supports. One hypothesis is that V1 comprises an array of neurons representing high-dimensional, non-linear random basis functions, capable of supporting a diverse set of downstream functions [Rahimi and Recht, 2007, Rahimi and Recht, 2008a, Rahimi and Recht, 2008b]. In addition to our results that showed that random-weight ReLU networks exhibit orientation-selective neurons, previous research on biologically plausible recurrent models of mouse V1 demonstrated the emergence of orientation selectivity even when both feedforward and recurrent connections are randomly initialized [Hansel and van Vreeswijk, 2012, Pattadkal et al., 2018].

Our findings contribute to the growing body of literature that emphasizes the importance of conducting controlled experiments to systematically investigate the architectural and training components that contribute to the neural encoding performance of computational models. Moreover, our results underscore the necessity of developing comprehensive batteries of neural and perceptual metrics to facilitate more informed conclusions about the similarities between computational models and the brain [Biscione et al., 2024, Jacob et al., 2021]. Finally, considering the computational benefits of the models’ representations that support the prediction of brain responses is valuable, as it helps formulate hypotheses regarding the functional roles of different brain areas [Dwivedi et al., 2021, Cichy and Kaiser, 2019].

## Methods

### Datasets

#### V1 monkey dataset

We used a public dataset that consists of neural activity recordings from 166 neurons across different layers of V1 brain area in two monkeys [Cadena et al., 2019]. The monkeys were shown 7,250 images, each presented 1-4 times for a duration of 60 milliseconds. Each image was displayed within a circular window spanning 2 degrees of visual angle, with the edges gradually fading out to blend with the surroundings.

#### IT monkey dataset

We used a publicly available IT monkey dataset which consists of neural recordings from 168 multiunit sites within the inferotemporal (IT) cortex of two macaque monkeys [Cadieu et al., 2014]. The monkeys were presented with 3,200 unique grayscale images, each showing one of 64 objects from eight categories. These images were designed to mimic real-world visual scenes by placing the cropped object images onto various natural image backgrounds at different positions, orientations, and sizes.

#### fMRI human dataset

The Natural Scenes Dataset (NSD) is a publicly available fMRI dataset that captures the brain activity of eight human participants as they viewed thousands of natural images (9,000–10,000 distinct color natural images for each subject repeated up to 3 trials) [Allen et al., 2022]. The images were taken from the Microsoft Common Objects in Context (COCO) database square-cropped and presented at a size of 8.4° x 8.4°. We used the regions of interest (ROI) V1v and V1d manually drawn based on the results of a population receptive field (pRF) experiment, and the higher-order ROIs VO1 and VO2, defined using a visual probabilistic atlas.

#### Texture-MNIST dataset

We created the Texture-MNIST dataset to probe different models’ texture and shape discrimination abilities. We created binary masks from the MNIST dataset, resized and overlaid them over 64 × 64 patches of texture randomly copped from a high-resolution texture image unique for each digit class. Using this dataset, we can train our models to either predict the class of the object (digit) or the class of the texture of each image.

### Models

We used simple DNN models consisting of a convolutional block and a linear readout. The convolutional block included two and four convolutional layers for the shallow and deeper models respectively. Each convolutional layer is followed by a batch normalization layer and an activation function. To maintain an efficient number of trainable parameters we used depthwise separable convolutions in all convolutional layers except for the first one [Du et al., 2024]. Furthermore, shallow models had 16 and 256 feature maps in their 2 convolutional layers, whereas deeper models had feature maps that progressively increased from 16 to 32, 64, and finally 256 across the network depth. For the shallow models, we had a pooling layer after each activation function and for the deeper models, we had the pooling layer after every other activation layer. For the shallow models, convolutional layers had either filter size of 9 × 9 or 17 × 17 pixels, leading to an effective receptive field of the models of 28 × 28 or 52 × 52 respectively. For the deeper models, we had filter sizes of 5 × 5 or 9 × 9 pixels leading to the same effective receptive fields as the sallow models (see supplementary Fig. S1 for an illustration of the detailed architecture of the shallow and deep 28 × 28 models). We tested a variety of activation functions including ReLU, ELU, Tanh, and Linear. Moreover, we considered average and maximum pooling operations, each with a pooling window of 2 × 2 pixels.

The three-dimensional activation maps of the convolutional block were transformed to the neural responses through a linear readout factorized using three one-dimensional weight vectors **w***_c_*, **w***_x_*, and **w***_y_* for the channels, and two spatial dimensions respectively. For the image classification tasks, global average pooling was applied to the three-dimensional activation maps to obtain the feature vector used for classification.

### Complexity measurement

To calculate the complexity of the function represented by a neural network, we evaluate the network on a regularly sampled grid in its input space [Teney et al., 2024]. Our networks were trained with input normalized to the range from −1 to 1. Therefore, we sampled 100 corners in the hypercube [−1, 1]*^d^*, where *d* is the input dimension of the network. We sampled 50 points regularly on each of the 100 lines connecting each corner with its succeeding corner and evaluated the network at each sample input point. Let *x* be the regularly sampled line in the range [−1, 1], and *y* be the activation of a certain output node evaluated at the data points lying on that line in the input space hypercube. We compute the coefficients **c** of Chebyshev polynomials that fit the data (*x, y*) by minimizing the least square error:

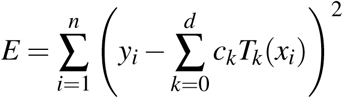

where *k* is the polynomial order and *T_k_*(*x_i_*) is the *k*−th Chebyshev polynomial evaluated at the *i*-th input *x_i_*. The Chebyshev polynomials *T_k_*(*x*) are recursively defined as:

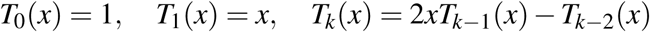

We define the complexity metric as the average of the polynomial orders weighted by their corresponding Chebyshev coefficients:

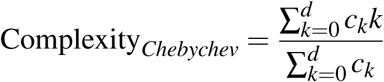

For each output node, we average the 100 complexity metrics from the 100 lines to obtain one complexity measurement per one output node.

### Comparison to experimental V1 data

To go beyond regression, we tested the alignment of the artificial neurons in our models with experimental data recorded from the V1 area by comparing their orientation selectivity distributions. We presented the models with Gabor patches of different orientations and phases [Kong et al., 2022]. Orientations were sampled at 10^◦^ steps in the range from 0^◦^ to 180^◦^ and 20 phases were sampled evenly from 0^◦^ to 360^◦^. The spatial frequency of the Gabor patches was chosen to allow the receptive field of the model neurons to contain two cycles i.e. for the receptive field of 28 × 28 pixels and input size of 128 × 128 pixels, we used a spatial frequency of 9.142 cycles/image. Specifically, we generated the Gabor patches according to the formula:

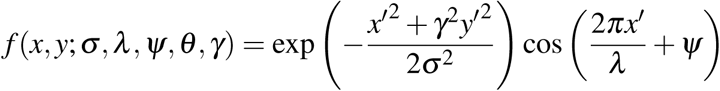

and

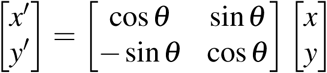

where *σ* = 20 pixels is the standard deviation of the Gaussian envelope which controls the size of the Gabor patch, *λ* is the wavelength of the sinusoidal factor in pixels, *ψ* is the phase offset of the sinusoidal factor, *θ* is the orientation of the Gabor patch in radians and *γ* = 1 is the spatial aspect ratio, specifying the ellipticity of the support of the Gabor function.

We calculated the orientation tuning curves of the center location of every artificial neuron at the last convolutional layer of our models. The tuning curves were scaled to the range from 0 to 1 to avoid negative responses in activation functions such as Tanh, Elu, and Linear. To establish a baseline for artificial neuron responses, the minimum response value across the V1 test set was determined. This minimum value was then used in the scaling of the tuning curves. Then we followed the analysis steps mentioned in [Gur et al., 2005] to calculate the orientation selectivity distribution of the V1 neurons recorded from awake monkeys. Briefly, we linearly interpolated the tuning curves with 1^◦^ steps, then smoothed them with a Hanning filter with a 7^◦^ half-width at half-height. We then quantified the orientation selectivity of neurons from the tuning curves by calculating the circular variance (CV) [Mazurek et al., 2014]. The circular variance was calculated from the smoothed tuning curves resampled at regular 15^◦^ intervals according to the equation:

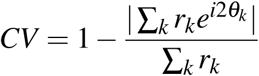

where *θ_k_* is the orientation in radians and *r_k_* is the corresponding response.

For each model, we simulated 100 in-silico electrophysiology experiments by randomly sampling with replacement 339 neurons from its last convolutional layer. We calculated their CV as described. From each experiment, we compared the distribution of CV with the corresponding distribution obtained from 339 V1 neurons recorded from alert macaques (Figure 3 in [Gur et al., 2005]) by calculating a V1 deviation score and then averaging the scores over the 100 experiments to obtain one score per model. The V1 deviation score was computed as the Wasserstein distance between the distribution of circular variance of the model’s neurons and the corresponding distribution of experimental V1 neurons.

## Acknowledgments

This project was financed by the BMF (Bundesministerium fuer Bildung und Forschung), Computational Life Sciences, project BINDA (031L0167); an ERC starting grant (850861) SPATEMP; DFG VI Grants (908/5-1 and 908/7-1); an NWO VIDI Grant; and the Dutch Brain Interface Initiative (DBI2).

## Supplementary Figures

**Figure S1.**
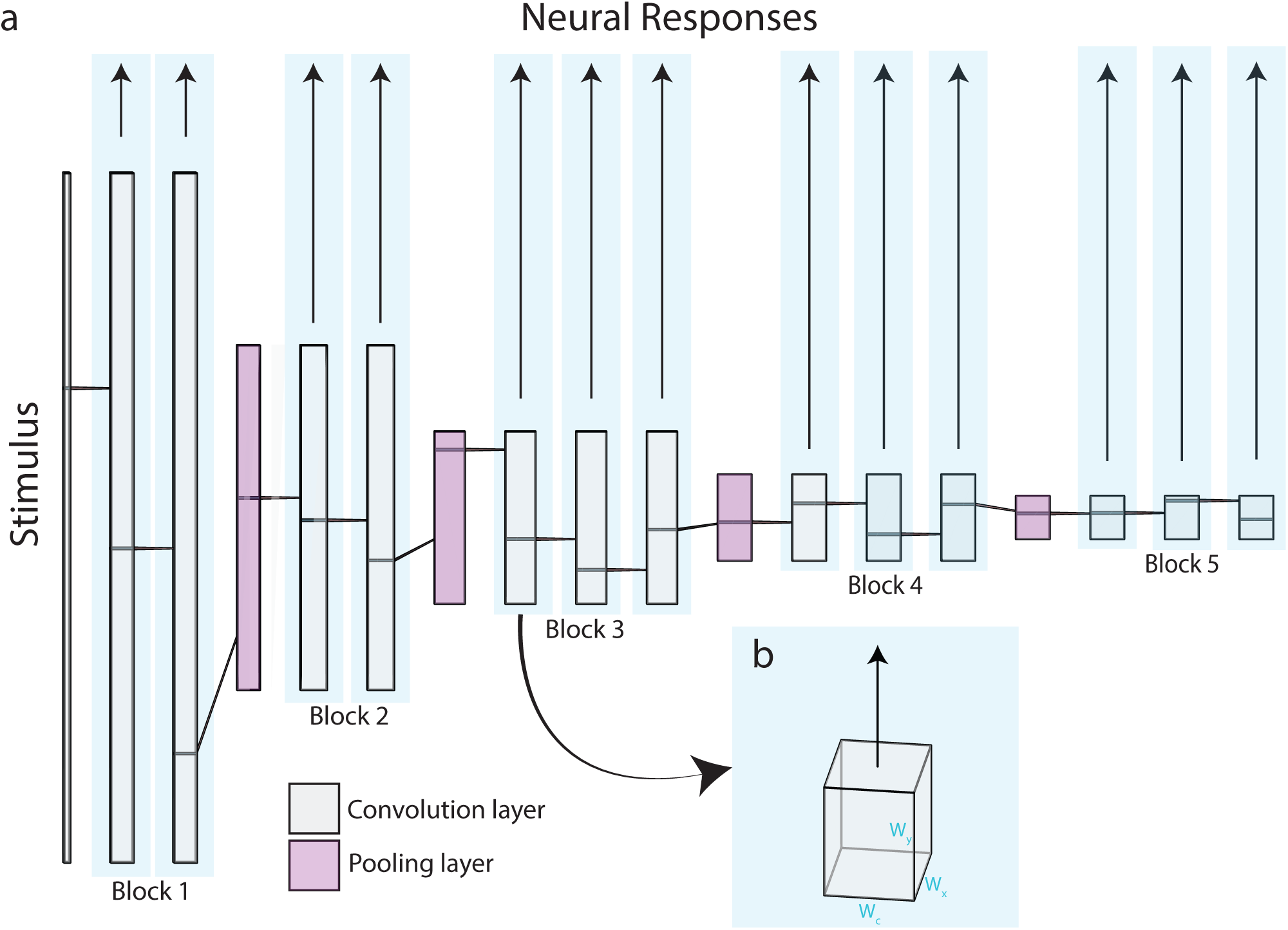
VGG16 encoding models. **(a)** On top of the three-dimensional feature maps of each convolutional layer (gray rectangles), we trained a linear readout (arrows) to predict the neural responses of a certain brain area. We used ImageNet-trained VGG16 model and randomly-initialized variants. **(b)** Linear readout was factorized into 3 one-dimensional weight vectors **w***_c_*, **w***_x_*, and **w***_y_*for the channels and the two spatial dimensions respectively.

**Figure S2.**
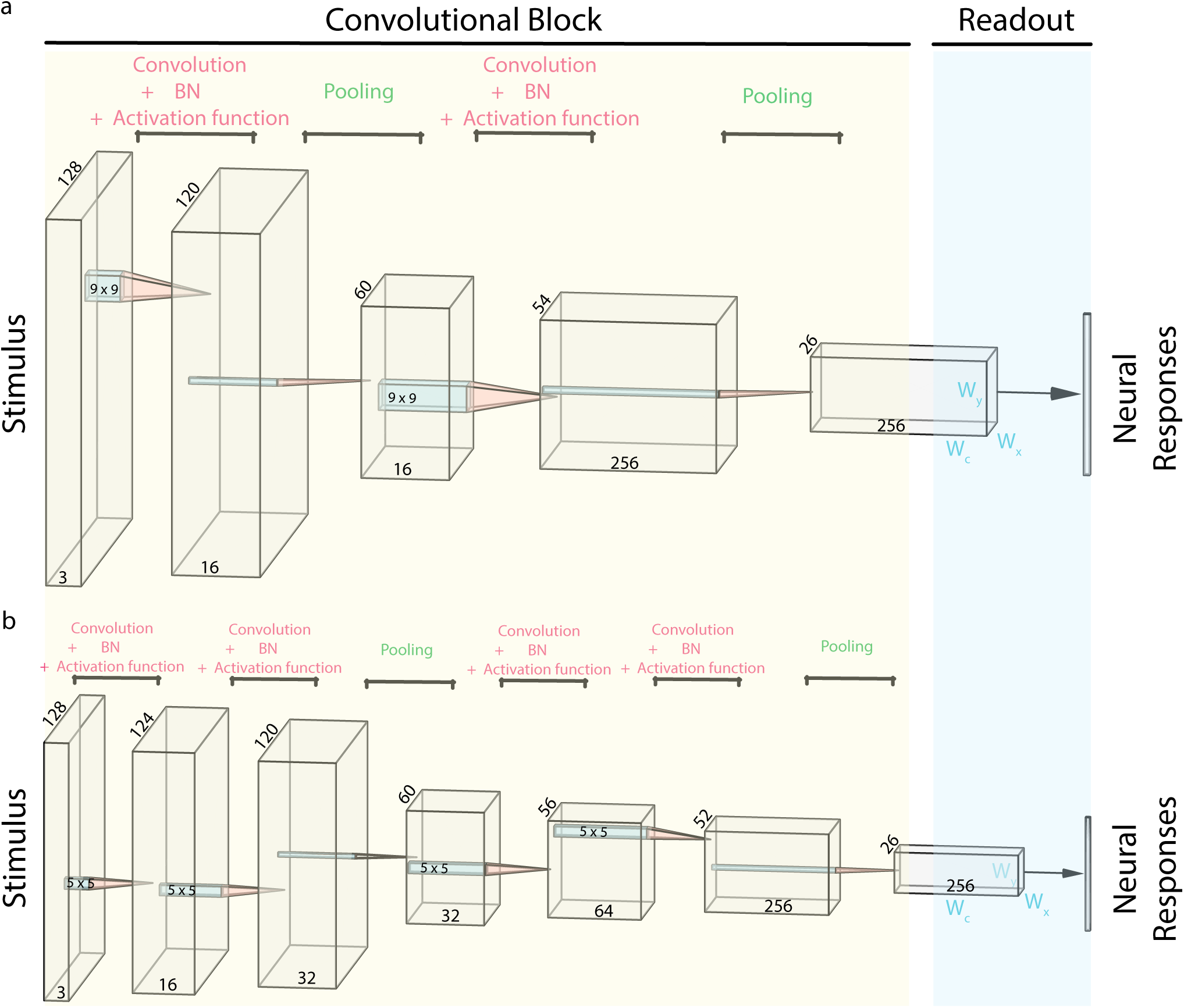
Architecture of CNNs used for encoding neural responses. All models are formed of a convolutional block and a linear readout. Readout is factorized into three one-dimensional vectors (**w***_c_*, **w***_x_*, and **w***_y_*) that transform the channels and the two spatial dimensions of the feature maps of the last convolutional layer into neural activity. **(a)** The convolutional block of shallow models used for encoding early visual cortex activity is formed of two convolutional layers with 9 × 9 filter sizes. Each layer is followed by a batch normalization (BN) layer, an activation function (ReLU, ELU, Tanh, or linear), and a pooling layer of 2 × 2 window size and stride = 2 (maximum or average pooling). **(b)** The convolutional block of the deeper models used for encoding higher visual areas is formed of 4 convolutional layers with 5 × 5 filter sizes. Each layer is followed by a batch normalization (BN) layer and an activation function (ReLU, ELU, Tanh, or linear). Every other layer is followed by a pooling layer of 2 × 2 window size and stride = 2 (maximum or average pooling). The spatial resolution of feature maps is printed on top of each block of feature maps. Number of feature maps (channels) is printed at the bottom. All convolutional layers are depthwise separable except for the first one. The effective receptive field of neurons in the feature maps of the last convolutional layer of both models is 28 × 28 pixels.

**Figure S3.**
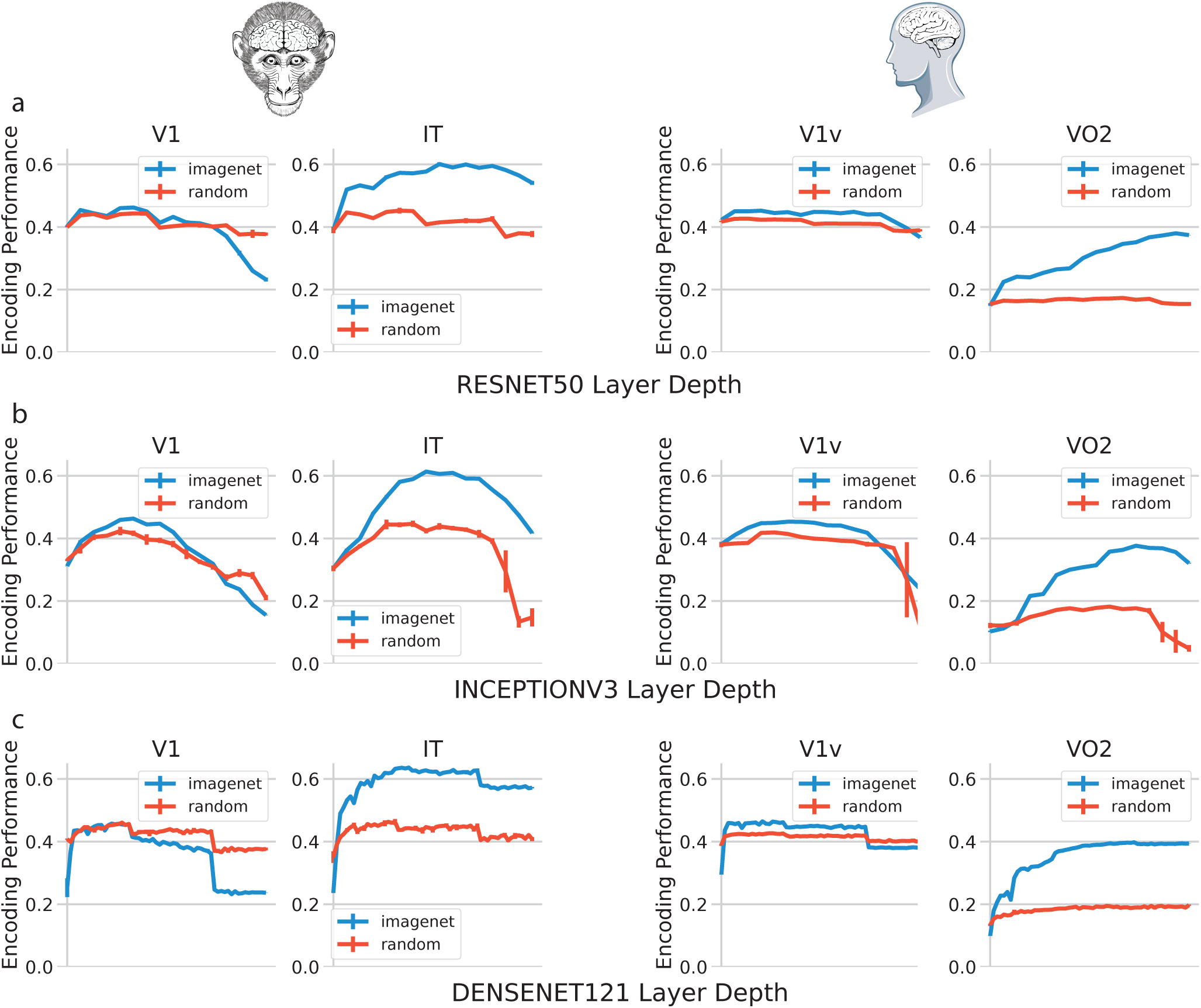
Encoding performance of popular convolutional architectures. **(a)** Encoding performance of a linear readout optimized on top of the representations of the convolutional layers of RESNET50 model either upon random initialization (in red) or pretraiend on ImageNet dataset for object recognition (in blue) for four neural datasets: Two electrophysiological datasets recorded from macaques: the early visual cortex V1 and the higher visual area IT and two fMRI datasets recorded from humans: the early visual cortex V1v and the higher visual area VO2. **(b)** same as **a** but for INCEPTIONV3 model. **(c)** same as **a** but for DENSENET121 model.

**Figure S4.**
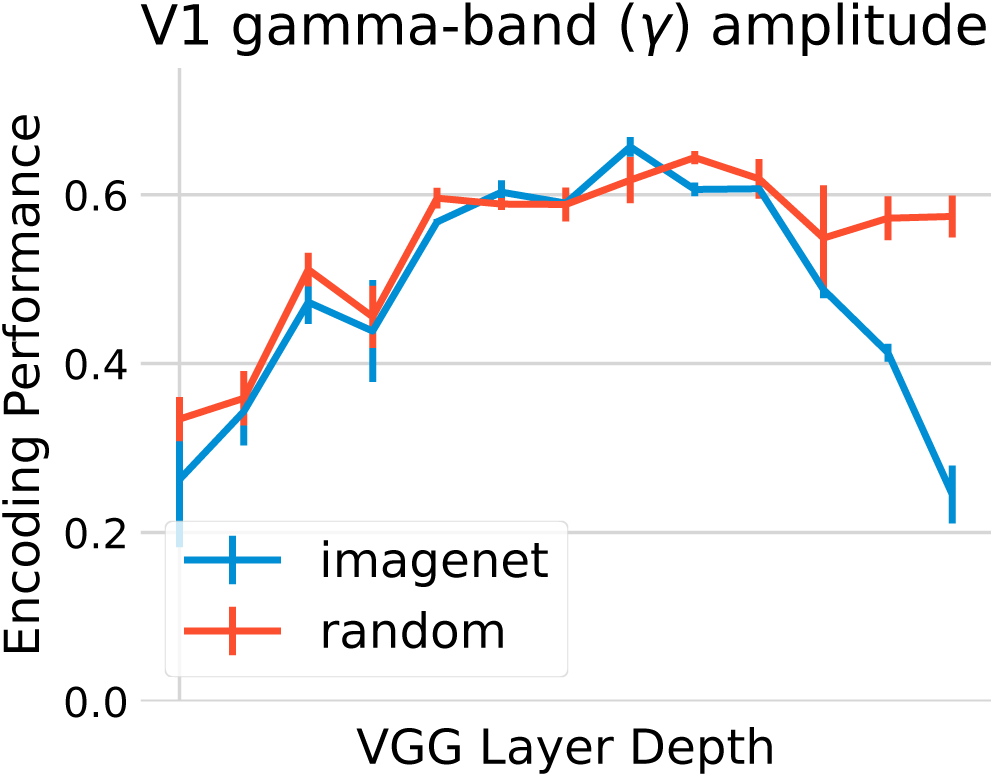
Data from [Uran et al., 2022]. In brief, recordings were made from one awake monkey passively viewing 11 degrees black/white natural images using multi-site recordings. Gamma-band local field potential (LFP) activity, which indexes rhythmic, synchronized activity in the local network, was measured by detrending the power spectrum and measuring the magnitude of the local spectral peak in the LFP. Note that gamma-band responses reflect surround modulations beyond the classic receptive field, which may explain the enhanced reliance on deeper layer representations for predicting V1 gamma-band activity [Uran et al., 2022, Peter et al., 2019, Vinck and Bosman, 2016, Gieselmann and Thiele, 2008].

**Figure S5.**
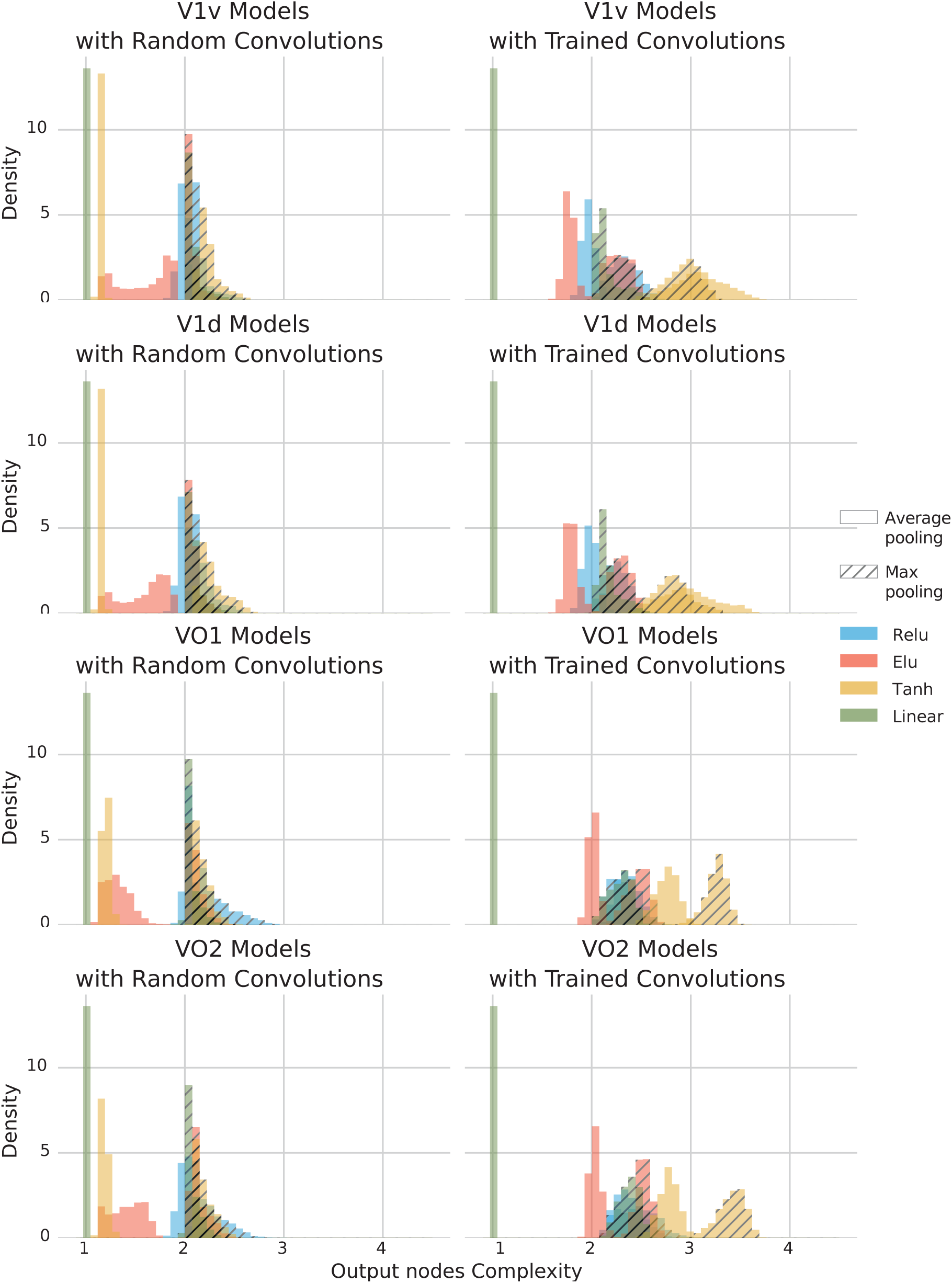
Distributions of encoding models’ output nodes complexity. Each distribution represent the complexity of output nodes (see methods) of a certain model configuration (with respect to the pooling strategy and activation function) trained on human V1v, V1d (upper two rows), VO1 and VO2 (lower two rows) data. Only linear readout was trained on top of random convolutional features (left column) or the model was fully trained (right column). Each distribution is the average of 3 training iterations.

**Figure S6.**
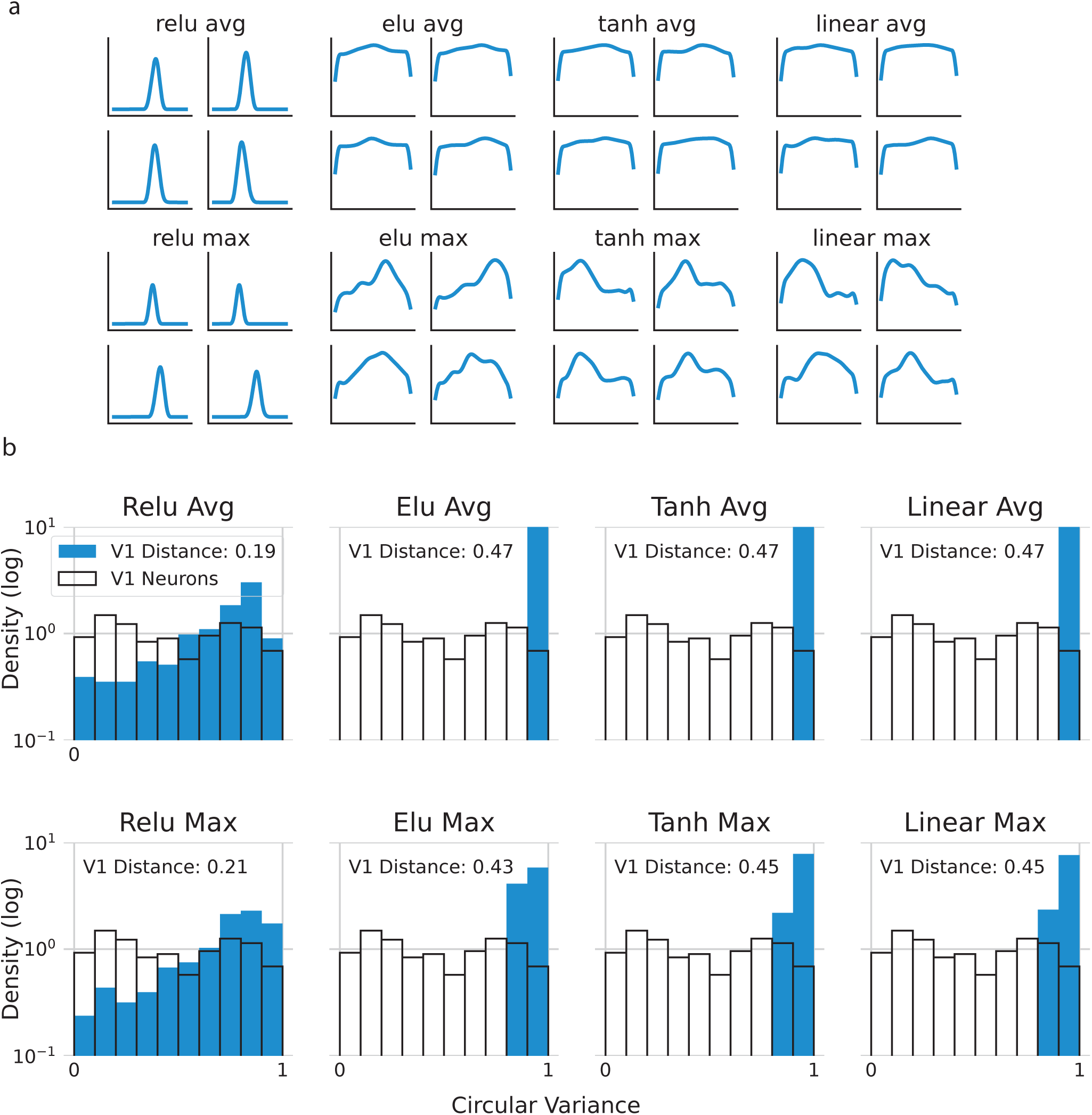
Orientation selectivity of random models. **(a)** Tuning curves of the most orientation-selective artificial neuron in the last convolutional layer of each model configuration upon random initialization. **(b)** Distributions of the circular variance of the artificial neurons of the last convolutional layer of each model configuration upon random initialization.

**Figure S7.**
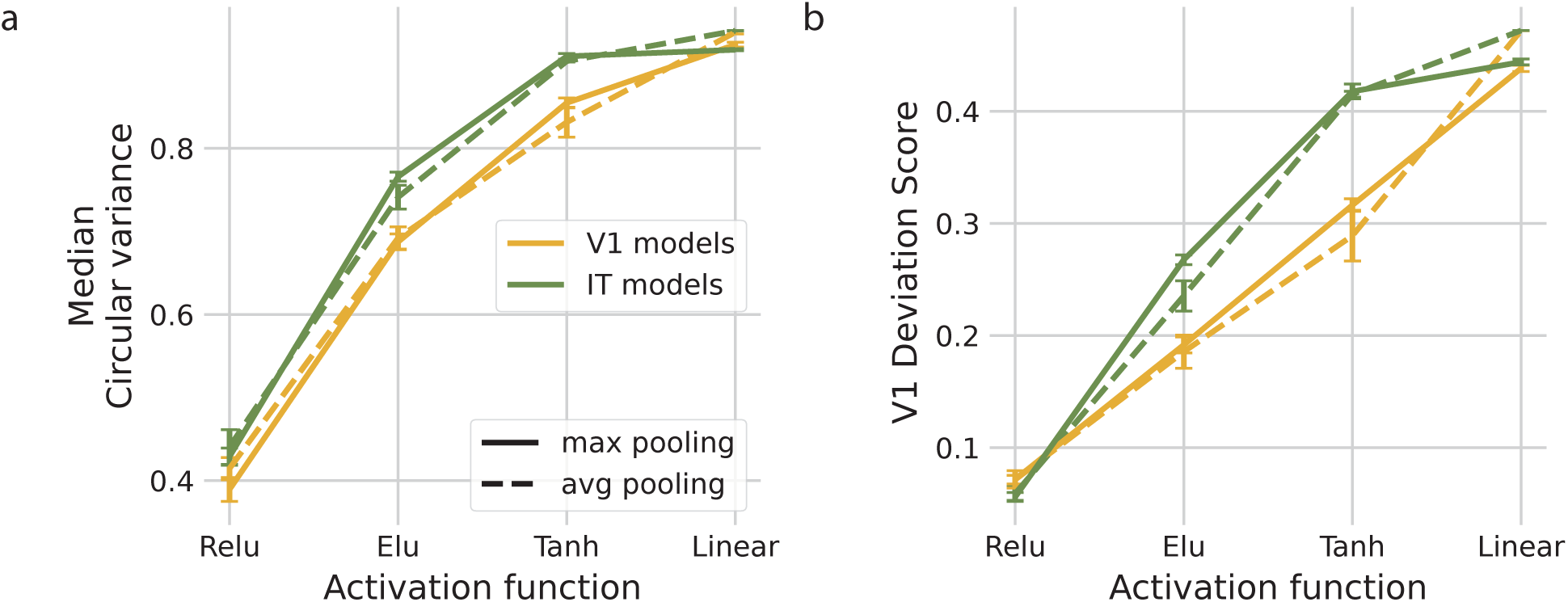
**(a)** Median circular variance of the artificial neurons of the last convolutional layer of V1 trained and IT trained models of different configurations averaged across 5 iterations. Error bars are the standard deviation. **(b)** V1 deviation scores of V1 trained and IT trained models averaged across 5 iterations. Error bars are the standard deviation.

